# Demonstration and structural basis of a therapeutic DNA aptamer for SARS-CoV-2 spike protein detection

**DOI:** 10.1101/2025.03.14.643408

**Authors:** Yujun Liu, Kaidong Wang, Weiguang Wang, Saarang Kashyap, Jonathan Jih, Anthony Imani, Tzung Hsiai, Z. Hong Zhou

**Affiliations:** California NanoSystems Institute, University of California at Los Angeles, Los Angeles, 90095, California, USA; Department of Microbiology, Immunology and Molecular Genetics, University of California at Los Angeles, Los Angeles, 90095, California, USA; Division of Cardiology, Department of Medicine, David Geffen School of Medicine, University of California Los Angeles, Los Angeles, California 90095, USA; Department of Medicine, Greater Los Angeles VA Healthcare System, Los Angeles, California 90073, USA; Molecular Biology Institute, University of California, Los Angeles (UCLA); Los Angeles, CA 90095, USA; Department of Bioengineering, Henry Samueli School of Engineering and Applied Science, University of California Los Angeles, Los Angeles, California 90095, USA

**Keywords:** Aptamer, cryo-EM, SARS-CoV-2, receptor binding domain, detection, specificity

## Abstract

At the onset of the COVID-19 pandemic, the absence of a rapid and highly specific diagnostic method for the SARS-CoV-2 virus led to significant delays in detection, adversely affecting clinical outcomes. This shortfall highlights the urgent need for adaptable, scalable, and reusable diagnostic technologies to improve future pandemic responses. To address this challenge, we developed a renewable electrochemical impedance biosensor device employing a synthetic nucleotide-based therapeutic aptamer (termed ‘aptasensor’) targeting the SARS-CoV-2 spike (S) protein receptor-binding domain (RBD). We demonstrate that our aptasensor can detect the Omicron BA.2 S protein within one hour and possesses concentration-dependent sensitivity at biologically relevant levels. Notably, the aptasensor is reusable after regeneration by a simple pH 2 buffer treatment. Aptamer binding to the S protein was confirmed by immunogold labeling and visualization by negative-stain electron microscopy. We used cryogenic electron microscopy (cryo-EM) to resolve high-resolution maps of the S protein in both the open and closed conformations and characterized aptamer binding to the up RBD in the open conformation. Taken together, these results establish the versatility and scalability of aptamer-based biosensors, presenting them as a potential transformative diagnostic platform for emerging pathogens. This combination of rapid detection, specificity, and renewable capabilities in a single diagnostic solution marks a significant advance in pandemic preparedness.

## 1. Introduction

The SARS-CoV-2 virus, a single-stranded positive-sense RNA virus of the *Coronaviridae* family, encodes four structural proteins in its genome: spike (S), envelope (E), membrane (M), and nucleocapsid (N) (Hoffmann et al. 2020; Walls et al. 2020). Among these, the S protein plays a central role in viral entry into host cells. It assembles as a homotrimer, with each monomer consisting of two functional subunits: S1, which binds to receptors, and S2, which mediates membrane fusion (Cai et al. 2020; Wrapp et al. 2020). The receptor-binding domain (RBD) within the S1 subunit specifically binds to the human angiotensin-converting enzyme 2 (hACE2) receptor and alternates between “open” and “closed” conformations (Shang et al. 2020), balancing receptor engagement with immune evasion. Following RBD binding to hACE2, the S2 subunit undergoes proteolytic activation by host proteases, exposing the fusion peptide required for facilitating membrane fusion between virus and host cell (Benton et al. 2020; Xia et al. 2020; Yan et al. 2020). These processes underscore the importance of the S protein in viral pathogenesis and its role as a key target for neutralizing antibodies and vaccine development.

The COVID-19 pandemic has emphasized the need for scalable and accurate detection methods to manage viral outbreaks. Current diagnostic approaches include molecular biological detection (e.g., reverse-transcriptase polymerase chain reaction, RT-PCR), immunological assays (e.g., enzyme-linked immunosorbent assays, ELISA), and pulmonary imaging techniques. RT-PCR, the gold standard for molecular detection, provides high sensitivity and specificity in identifying viral RNA but is limited by its dependence on sophisticated infrastructure and susceptibility to mutations in viral target sequences (Artika et al. 2022; Samson et al. 2020).

Immunological assays detect viral proteins more rapidly but require specialized equipment and expertise (Khan et al. 2022), while pulmonary imaging methods, such as chest X-rays or computed tomography, are labor-intensive and cost-prohibitive at scale (Shi et al. 2020). These limitations highlight the urgent need for diagnostic technologies that are rapid, accessible, and adaptable to evolving viral threats.

Biosensors represent a promising alternative to traditional diagnostic methods. Such sensors convert biological signals into measurable electrical outputs through optical, electrochemical, piezoelectric, or thermal modalities (Justino et al. 2016; Perumal and Hashim 2014). These devices offer advantages such as high sensitivity, rapid detection, and potential scalability for point-of-care applications (Pohanka 2018). Their widespread adoption, however, is hindered by challenges such as short operational lifespan, limited reusability, and dependence on multiple devices for signal detection and processing (Patel et al. 2023). Overcoming these obstacles is crucial to unlocking the full potential of biosensors for infectious disease diagnostics.

Aptamers have emerged as a transformative solution for biosensor development. These single-stranded nucleic acids, selected through systematic evolution of ligands by exponential enrichment (SELEX), exhibit high-affinity and high-specificity binding to target molecules comparable to antibodies (Sefah et al. 2010). They offer distinct advantages, however, including lower production costs, enhanced chemical stability, and ease of modification (Kumar Kulabhusan et al. 2020). Recent breakthroughs in machine learning have further refined the design and synthesis of aptamers, enabling rapid adaptation to mutations in viral targets (Bashir et al. 2021; Chen et al. 2021). These advancements position aptamers as versatile and scalable solutions for next-generation biosensors, addressing key limitations of traditional diagnostic methods while providing the adaptability needed to counter emergent threats effectively.

Among emergent threats, SARS-CoV-2 stands out as a highly mutagenic pathogen that necessitates rapid and precise diagnostic solutions. In this study, we developed a renewable aptamer-based electrochemical impedance biosensor device, termed an aptasensor, to detect the SARS-CoV-2 S protein. Our aptasensor, composed of a thiol-functionalized anti-SARS-CoV-2 S protein aptamer immobilized on gold nanoparticle-modified carbon nanotube electrodes, achieved rapid and dose-dependent detection of the Omicron BA.2 S protein within a short timeframe. To further characterize native aptamer binding, we used cryo-EM to confirm its selective interaction with the open conformation of the S protein and observed no binding in the closed state. Importantly, the aptasensor retained its functionality after simple pH-based regeneration, underscoring its reusability. This innovative design highlights the potential of the aptasensor as a scalable, reusable, and adaptable diagnostic platform for SARS-CoV-2 and other emergent viral threats.

## 2. Materials and methods

### 2.1 Reagents

Gold (III) chloride hydrate (∼52% Au basis), potassium chloride hydrate, sulfuric acid (95.0-98.0%), potassium hexacyanoferrate (II) trihydrate (≥98.5%), potassium ferricyanide (III) (99%), HEPES solution, Tween 20, glycine (≥98.5%), ethanol (≥99.5%), sodium chloride (≥99.0%), phosphate buffer solution (PBS, 1.0 M, pH 7.4), and bovine serum albumin (BSA) were obtained from Sigma-Aldrich. All reagents were chosen for their high purity to ensure consistency and reproducibility in sensor fabrication. Multi-walled carbon nanotubes (XFM01, 5-15 nm in diameter, 10-30 μm in length) were sourced from XFNANO Technology Co. Ltd. The anti-SARS-CoV-2 S protein aptamer CoV2-6C3 (5’-CGC AGC ACC CAA GAA CAA GGA CTG CTT AGG ATT GCG ATA GGT TCG G -3’) (Sun et al. 2021) was synthesized and purified by Integrated DNA Technologies (IDT). Recombinant SARS-CoV-2 S protein (lyophilized and liquid forms) was purchased from ACROBiosystems (SPN-C5227, Newark, DE, USA).

Ultrapure water (18.2 MΩ cm) was prepared using a Millipore purification system.

### 2.2 Electrochemical impedance tests

Electrochemical measurements were conducted using a Gamry Interface 1010E workstation (Gamry Instruments, Inc., USA). Electrochemical impedance spectroscopy (EIS) was performed over a frequency range from 100 kHz to 0.1 Hz to capture relevant charge transfer dynamics, using an electrolyte solution containing 0.1 M KCl, and 5 mM [Fe (CN)_6_]^3-/4-^. This composition was selected to facilitate electron transfer between the electrode and the analyte. The Randles model was used to fit the EIS data, with charge transfer resistance (*R*_ct_) values extracted to evaluate aptamer binding events and sensor performance (Vivier and Orazem 2022).

### 2.3 Preparation of a label-free electrochemical impedance aptasensor

*Carbon Nanotube Dispersion and Electrode Preparation* A 2 mg/mL suspension of multi-walled carbon nanotubes (CNTs) in a 50:50 (v/v) water-ethanol mixture was sonicated for 2 hours to ensure uniform dispersion and prevent aggregation. A glassy carbon electrode (GCE, 3 mm diameter) was polished with 50 nm aqueous alumina slurry on a polishing cloth to enhance surface roughness and cleaned via ultrasonication in ethanol and water. The cleaned GCE was dried under a nitrogen gas stream to avoid contamination.

*CNT Deposition and Gold Nanoparticle (AuNP) Coating* 5 μL of the CNT suspension was drop-cast onto the GCE surface and dried at room temperature to form a CNT-modified GCE (CNTs/GCE). Gold nanoparticles (AuNPs) were electrochemically deposited onto the CNTs/GCE by cycling the potential in a solution of gold (III) chloride hydrate solution in 0.5 M H_2_SO_4_. The resulting AuNPs/CNTs/GCE served as the substrate for aptamer immobilization, with AuNPs enhancing conductivity and providing a binding platform for thiolated aptamers.

*Aptamer Immobilization and Blocking* A thiol-functionalized anti-SARS-CoV-2 S protein aptamer solution was incubated on the AuNPs/CNTs/GCE for 1 hour at room temperature to form a self-assembled monolayer. After aptamer immobilization, the electrode was rinsed with phosphate-buffered saline (PBS, pH 7.4) to remove unbound aptamers. To prevent non-specific binding, the surface of the electrode was treated with 0.05% BSA solution in PBS for 30 minutes, followed by rinsing with PBS. The aptamer/AuNPs/CNTs/GCE electrochemical impedance aptasensor was stored at 4°C for further use.

### 2.4 Surface plasmon resonance

Binding kinetics were analyzed using a Biacore 2000 system at 25°C. The aptamer was immobilized onto a Series S Sensor Chip SA (GE Healthcare) in a running buffer containing 10 mM HEPES, 150 mM NaCl, pH 7.4, and 0.005% (v/v) Tween-20. Serial dilutions of the SARS-CoV-2 S protein were injected at a flow rate of 30 µL/min, with association and dissociation phases lasting 180 seconds and 900 seconds, respectively. Regeneration of the chip surface was achieved using 0.1 M glycine-HCl buffer (pH 2) applied in two 10-second intervals. The optimized buffer composition ensured aptamer stability and reproducible sensor performance across cycles.

### 2.5 Negative-stain sample preparation and microscopy

Aptamer-conjugated AuNPs were incubated with SARS-CoV-2 S protein on ice for 30 minutes at a 1:1 molar ratio. Glow-discharged Formvar-coated copper electron microscopy (EM) grids (Electron Microscopy Sciences, USA) were prepared by applying the sample for one minute, blotting excess liquid, then washing three times with 2% uranyl acetate. A final incubation with uranyl acetate for one minute provided contrast for imaging. Grids were dried at room temperature and imaged at 15,000x magnification using a Tecnai T12 microscope (Thermo Fisher Scientific, USA). Control samples, prepared using the same protocol without aptamers, were used to verify specificity.

### 2.6 Cryo-EM sample preparation and imaging

The aptamer was incubated with thawed SARS-CoV-2 S protein (500µg/dL) on ice for 30 minutes at a 4:1 molar ratio. A 4 µl aliquot of sample was applied to holey carbon Quantifoil R2/1 copper 200-mesh grids (Electron Microscopy Sciences, USA), blotted, and vitrified using a Vitrobot Mark IV (Thermo Fisher Scientific, USA) in a liquid ethane/propane mixture. Grids were screened in a Tecnai T20 (Thermo Fisher Scientific, USA) transmission electron microscope to optimize freezing conditions.

Cryo-EM data was collected on a 300 kV Titan Krios (Thermo Fisher Scientific, USA) equipped with K3 direct electron detector (Gatan, USA) operated in electron counting mode. Movies were recorded at 81,000× magnification with a dose rate of 26.97 e^-^ pixel^-1^ s^-1^. A total of 12,834 movies were captured, each comprising 40 frames with a total exposure time of 2 seconds. Automated image acquisition was accomplished using SerialEM (Mastronarde 2018).

### 2.7 Data processing and 3D reconstruction

Data processing for 3D reconstruction was processed using CryoSPARC v4.3.1 and RELION-4.1 (Punjani et al. 2017; Scheres 2012). Micrograph curation was performed to remove low-quality images before proceeding with particle picking. In total, 11,972 selected micrographs were subjected to patch motion correction using *MotionCor2* and CTF estimation using *GCTF* (Zhang 2016; Zheng et al. 2017).

A previously published closed conformation map EMD-30660 (Xu et al. 2021) was used as a reference to generate templates for automated picking using *Template Picker* in CryoSPARC. These automatically picked particles were then subjected to iterative 2D classification, where the best classes were selected as templates for a second round of particle picking and subsequent 2D classification. The best classes were used to train a Topaz model (Bepler et al. 2019) for improved S protein particle picking. The trained Topaz model picked 1,175,879 particles, which were extracted at a box size of 330 Å (300 pixels) and filtered through additional rounds of 2D classification to remove junk particles, yielding a final dataset of 671,570 particles.

One round of homogeneous refinement on the extracted particles showed poorly resolved RBD densities in the resulting map, indicating multiple S protein conformations being averaged together. To resolve these different map conformations, we subsequently performed 3D classification on the newly extracted particles and specified 10 different classes. All 10 classes were then subjected to heterogenous refinement with the previous 3D classification volumes as references, revealing two major classes corresponding to the open and closed conformation maps, respectively. The other classes from this heterogenous refinement step were poorly aligned or appeared to be missing spike protein density, so we decided to discard them. The two major classes were then subjected to another round of heterogeneous refinement to remove more bad quality particles contributing to the two major conformations maps. One round of non-uniform refinement (Punjani et al. 2020) in C1 symmetry was performed for both open and closed maps, followed by a C3 refinement for the closed conformation due to its symmetrical positioning of the down RBDs. This helped improve the resolution of the closed conformation map to 3.4 Å (Table S1).

To better resolve the aptamer-RBD interaction in the open conformation, additional data processing was carried out in RELION. A round of 3D classification was conducted on the entire structure to remove particles corresponding to closed and misaligned conformations, followed by a final round of global refinement. This refinement revealed a low-resolution helical density with distinct grooves binding to the top of the open confirmation up RBD. Since this density aligned well with our aptamer model, we confidently assigned it as the aptamer. To mitigate preferred orientation artifacts in the open conformation map, anisotropic and misalignment correction modules in *spIsoNet* (Liu et al. 2025) were applied. Sharpening and postprocessing were performed on the half-maps generated from spIsoNet, yielding a final open confirmation map at 4.4 Å resolution (Table S1).

To assess structural variability of the up RBD in the open conformation, 3D variability analysis (Punjani and Fleet 2021) was conducted in CryoSPARC, performed across three different modes with the “filter resolution (Å)” parameter set at 6 Å. A total of 20 volumes per mode was generated and volumes were downsampled to 150px, capturing dynamic structural changes within the dataset. These volumes were compiled into a movie to visualize motion and flexibility (Movie S1), providing deeper insights into conformational dynamics and potential interactions.

### 2.8 Atomic model building and refinement

Initial model building was performed in Coot-0.8.9 (Emsley et al. 2010) and ISOLDE (Croll 2018) using sharpened maps for both conformations: the open conformation model was adapted from PDB 7ND9 (Dejnirattisai et al. 2021) and the closed conformation model was derived from PDB 7DF3 (Xu et al. 2021). Models of the open conformation were flexibly fitted into our 3D variability maps to track positional changes of the up RBD. Model building of the aptamer structure was based on initial AlphaFold3 (Abramson et al. 2024) prediction of CoV2-6C3 sequence, which was further refined to ensure correct base pairing of the aptamer. Following initial model building, amino acid residues of the S protein and nucleotides of the aptamer were both fixed via positional restraints. However, RBD residues involved in nucleotide recognition and interaction, along with nucleotides predicted to interact with the RBD based on position and literature, were released from these constraints, along with their neighboring residues (Sun et al. 2021). Potential RBD-aptamer interactions were simulated by implementing appropriate distance restraints for noncovalent forces between the aforementioned amino acids and nucleotides. Our ISOLDE simulation allowed us to flip out nucleotide bases and flexibly model non-covalent interactions with the aptamer residues that conformed to our density map contact points. Final refinements were performed in ISOLDE to optimize interactions between the aptamer and the open RBD. Structure visualization and figure preparation were done with UCSF ChimeraX (Meng et al. 2023).

## 3. Results

### 3.1 Development and performance of an electrochemical aptasensor for SARS-CoV-2 detection

The global urgency for rapid, sensitive, and scalable diagnostic tools has driven advancements in biosensor technology. Towards this end, we set out to develop an electrochemical impedance aptamer-based biosensor device, or aptasensor, leveraging a therapeutic aptamer sequence that targets the RBD of the SARS-CoV-2 S protein. To maximize detection range of the aptasensor, carbon nanotubes (CNTs) were used as base electrodes due to their large specific surface area to better facilitate aptamer conjugation onto electrochemically deposited gold nanoparticles (AuNPs) (Fig. 1a). Electrochemical impedance spectroscopy (EIS) was used to monitor the charge-transfer resistance (*R*_ct_) values, modeled using the Randles equivalent circuit model (Fig. 1b) (Vivier and Orazem 2022). Nyquist plots (Fig. 1b) revealed a semicircular region indicative of charge-transfer limitation at high frequencies, and a linear portion corresponding to diffusion-limitation at low frequencies (Vivier and Orazem 2022). The *R*_ct_ values increased proportionally with SARS-CoV-2 S protein concentrations, demonstrating specific adsorption and subsequent charge-transfer impedance in the presence of the [Fe (CN)_6_]^3-/4-^ redox probe. The aptasensor achieved a detection range of 1 pg mL^-1^ to 10^5^ pg mL^-1^, with a strong linear correlation (*R*^2^ = 0.995) (Fig. 1c) and a limit of detection (LOD) of 0.19 pg mL^-1^ (S/N = 3). This performance reflects the synergistic impact of CNT-enhanced sensitivity and aptamer specificity (Wongkaew et al. 2018). Notably, the biologically significant serum S protein concentration, reported as 33.9 ±22.4 pg/mL (Yonker et al. 2023), falls within the detection range of the aptasensor. Moreover, the aptasensor detected SARS-CoV-2 S protein within one hour, highlighting its potential for real-time diagnostics. The ability of the aptasensor to detect SARS-CoV-2 S protein with high sensitivity and in a biologically relevant range underscores its practical utility.

**Figure 1.**
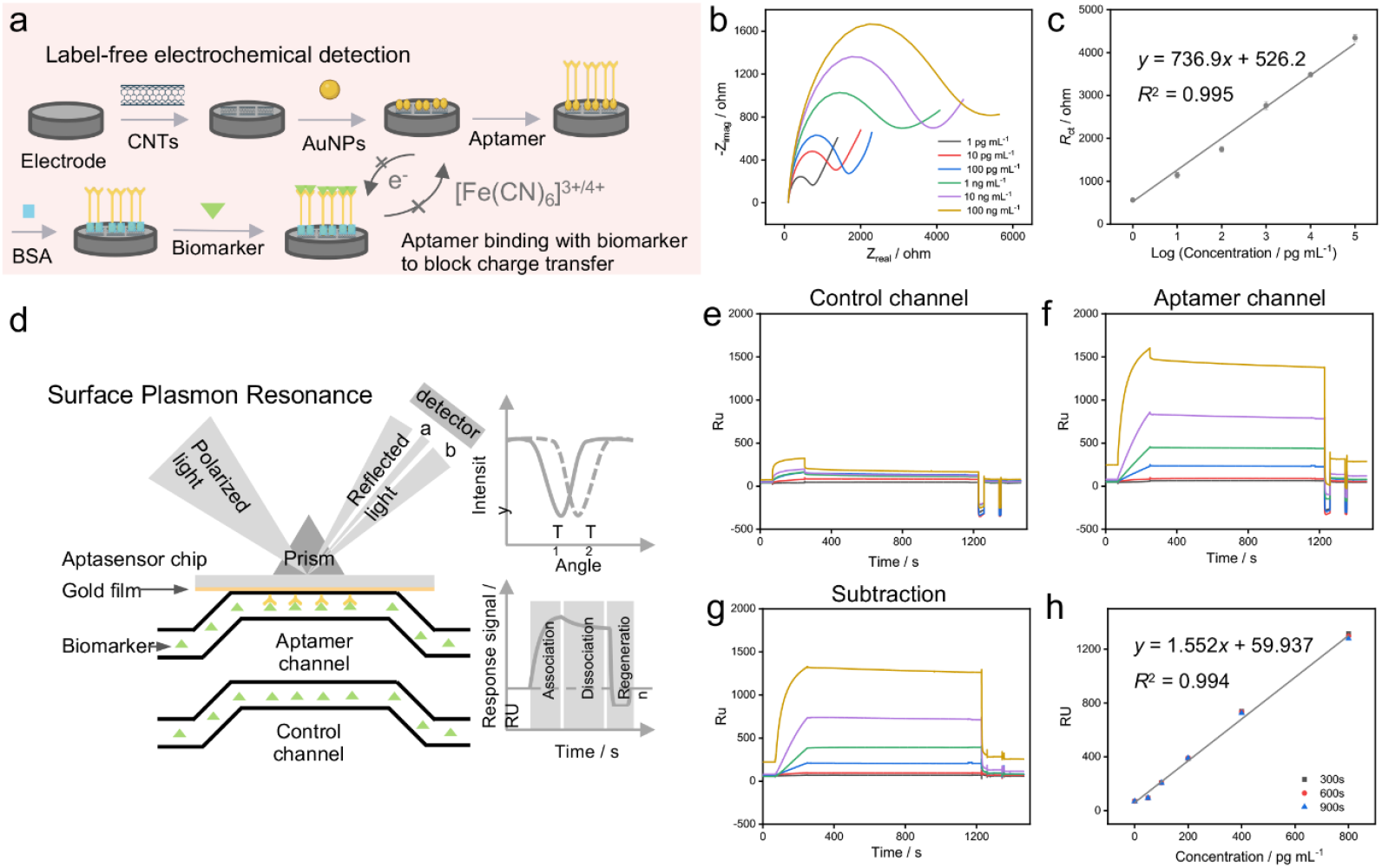
Development and performance of aptasensor for SARS-CoV-2 S protein detection. (**a**) Schematic representation of the electrochemical impedance aptasensor fabrication process using carbon nanotubes (CNTs), gold nanoparticles (AuNPs), and an anti-SARS-CoV-2 S protein aptamer for selective detection. (**b**) Electrochemical impedance spectroscopy (EIS) responses at different concentrations of SARS-CoV-2 S protein, ranging from 1 pg/mL to 10^5^ pg/mL. (**c**) Calibration curve for correlating EIS response (R_*ct*_ values) with SARS-CoV-2 S protein concentrations, fitted using the Randles equivalent circuit model. (**d**) Diagram illustrating the working principle of surface plasmon resonance (SPR) detection for SARS-CoV-2 S protein detection using an aptamer-modified microfluidic chip. (**e**) SPR signal response recorded in the control channel without aptamer modification, indicating nonspecific adsorption. (**f**) SPR signal response observed in the aptamer functionalized channel, demonstrating specific binding to the SARS-CoV-2 S protein. (**g**) Subtracted SPR signal response to highlight the specific detection of the SARS-CoV-2 S protein by eliminating background signals from the control channels. (**h**) Calibration curve for SPR responses at different concentrations of SARS-CoV-2 S protein.

To assess the binding kinetics and specificity of the aptamer, we employed surface plasmon resonance (SPR). The SPR response increased proportionally with SARS-CoV-2 S protein concentrations, confirming high-affinity binding between the aptamer and the S protein (Figs. 1 d-h) (Hanke et al. 2022). Regeneration studies demonstrated that the aptasensor retained binding efficiency across multiple cycles following treatment with a 0.1 M glycine-HCl buffer (pH 2). This reversible binding is attributed to non-covalent forces, such as hydrogen bonding and Van der Waals interactions (Williams et al. 2021), confirms the durability and cost-effectiveness of the aptasensor for repeated diagnostic use. These findings highlight its strong binding affinity, reusability, and potential as a scalable diagnostic platform for SARS-CoV-2 and emerging pathogens.

### 3.2 Cryo-EM reveals aptamer-S protein interactions

We used immunogold labeling to confirm the presence of aptamer-conjugated AuNPs and visualized their association with S protein via negative-stain EM (Fig. 2a). Initial cryo-EM studies using lyophilized S protein revealed that it had lost its ability to interact with the aptamer.

**Figure 2.**
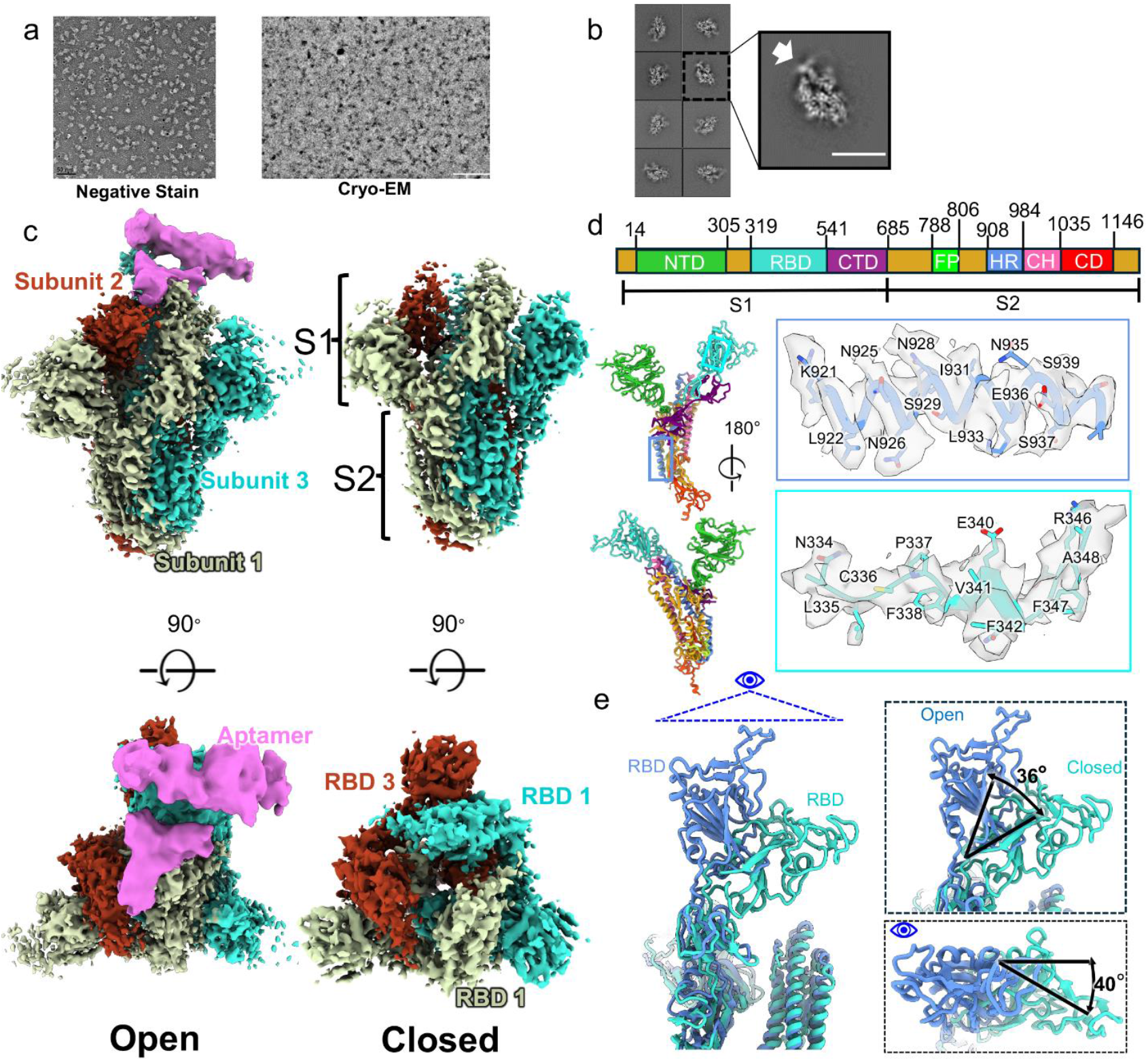
Structural analysis of aptamer binding to SARS-CoV-2 S protein trimer. (**a**) Representative micrographs of negative-stain microscopy of aptamer-conjugated gold nanoparticles bound to S protein (Scale bar: 50 nm) and of cryo-EM of aptamer-S protein complexes (Scale bar: 100 nm). (**b**) Representative 2D class averages of aptamer-S protein complexes in cryo-EM images, with aptamer density interfacing with the receptor-binding domain (RBD) indicated by a white arrow. Scale bar: 135 Å (**c**) Maps of the S protein in open and closed conformations, with the aptamer densities in pink. (**d**) Labeled domains of full-length SARS-CoV-2 S protein. The S1 subunit contains NTD, RBD, and CTD, while S2 contains FP, HR1, CH, and CD. (**e**) Comparison of RBD positioning between open and closed conformations.

To determine the structure of the aptamer-S protein complex, we recorded ∼12,000 movies using cryo-EM with liquid S protein and subjected our extracted S protein particles to single particle analysis (Fig. 2b). Extensive 3D classification and heterogenous refinement revealed two major classes with distinct differences in RBD positioning and aptamer interactions (Fig. 2c). We designated these classes as open and closed conformations consistent with previous studies (Ye et al. 2022). Further 3D refinement led to an open confirmation map at resolution 4.4 Å and a closed conformation map at 3.4 Å.

The open conformation of the S protein consists of one RBD in an upright position (up RBD) and two in the lying-down position (down RBD), with the aptamer bound to the up RBD. The aptamer binds with the up RBD at two distinct locations, forming a bridge-like interaction similar to the three-pointed hACE2 interaction model (Wrapp et al. 2020). Additionally, we observed a second sickle-shaped density that binds within the cleft between the two down RBDs and the one up RBD. In contrast, the closed conformation has all three RBDs in the down position with no residues exposed for aptamer binding.

To gain further insight into RBD–aptamer interactions and positioning, we atomically modeled the open and closed S protein conformations, assigning domains to our monomers based on established literature: N-terminal Domain (14-305), RBD (319-541), C-terminal Domain (541-685), Fusion Peptide (788-806), Heptapeptide Repeat Sequence 1 (908-984), Central Helix (984-1035) and Connector Domain (1035-1146) (Yan et al. 2020) (Fig. 2d). Our modeling of the S1 (residues 1–685) and S2 (residues 686–1164) subunits revealed that while the S2 domains remain relatively rigid, the RBDs within S1 exhibit greater flexibility (Movie S1). In the S1 domain, the up RBD is elevated ∼ 3.7 nm from the down RBD, with an approximate 36° angle separating the two RBD configurations and a 40° angle rotation out of the page (Fig. 2e).

These findings highlight the critical role of RBD movement and RBD exposure in aptamer recognition, providing a structural framework to optimize aptamer-based detection strategies.

### 3.3 Positional variability and interactions of aptamer-RBD complex

The open conformation of the S protein binds the aptamer through its up RBD, forming a complex that is nearly perpendicular to adjacent monomer NTDs. To characterize the flexibility of this aptamer-RBD complex, we employed 3D Variability Analysis (3DVAR) in CryoSPARC to model its movement relative to the rest of the S protein (Fig. 3a). Our results revealed that the aptamer-RBD complex shifted up to 2 nm towards the adjacent RBD, transitioning from a “straight” conformation to a “bent” conformation (Fig. 3b). Notably, the aptamer-RBD complex transiently shifts between the straight and bent conformations, showing strong binding to both conformations but reduced aptamer binding in the transition between them.

**Figure 3.**
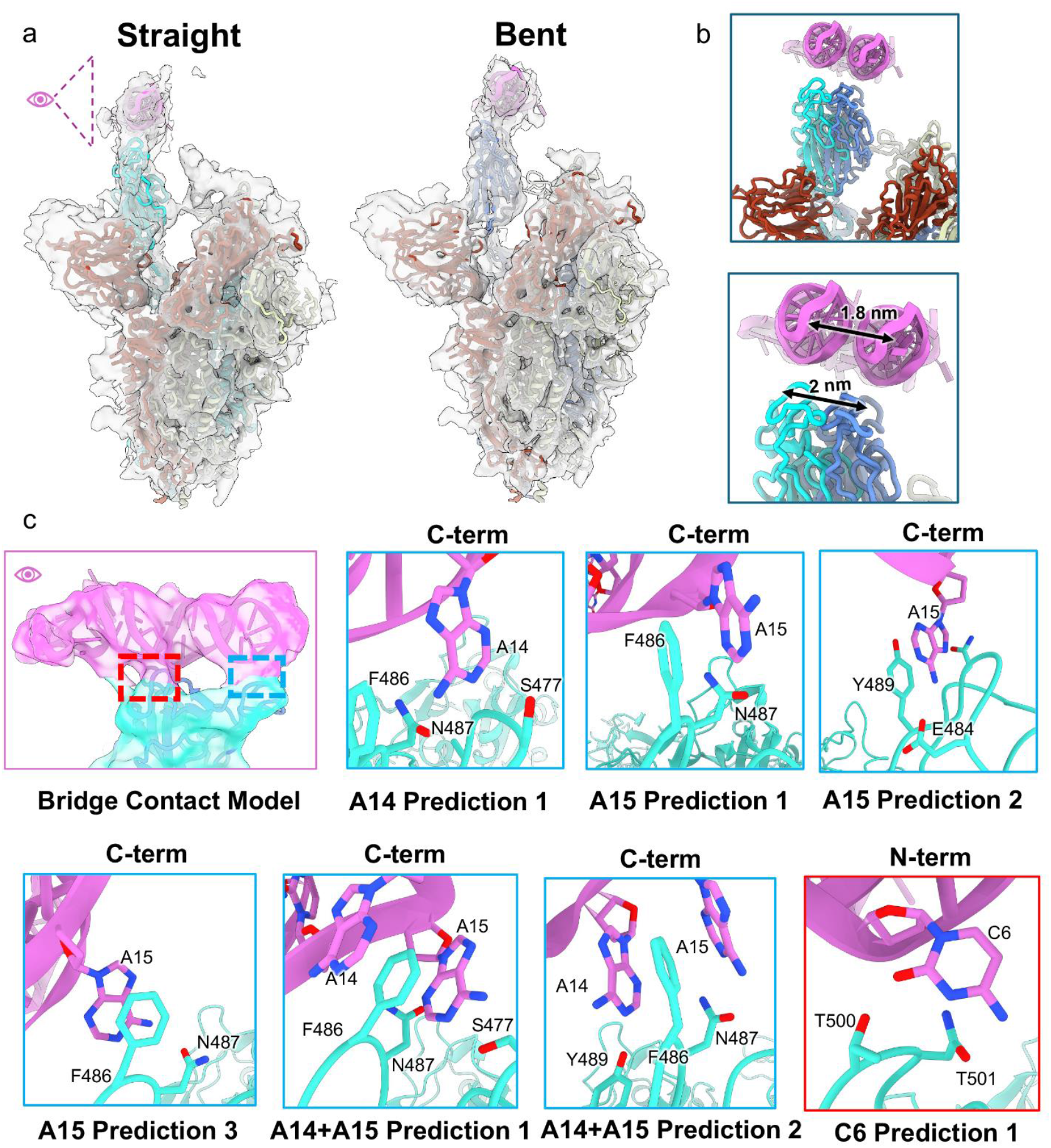
Flexibility of aptamer-RBD complex and density-guided simulations of aptamer binding interactions. (**a**) 3D Flexibility of the open conformation in the Straight and Bent positions. (**b**) shift in the aptamer-RBD position from straight to bent conformation. (**c**) Density-guided ISOLDE simulations of interactions between aptamer nucleotides and RBD residues at the C- and N-terminal regions of the RBD.

To further characterize the bridge-like interface between the aptamer and the RBD, we modeled putative interactions at the C-terminal region (residues 445-490) and the N-terminal region (residues 490-505) of the S protein RBD in the RBD-aptamer complex. Since our map precluded atomic modeling of interactions at the aptamer-RBD interface, we leveraged ISOLDE, a density-guided simulation software, to reveal potential binding mechanisms of the aptamer to the RBD. ISOLDE simulations steered the fitting of both S protein residues and aptamer nucleotides into the map, while considering noncovalent interactions (hydrogen bonds, ionic interactions, and van der Waals forces) and maintaining geometry constraints (bond lengths and angles) for protein and nucleic acid components. Based on our simulations, we propose that the aptamer interacts with the C-terminal cluster of RBD amino acids Y489, N487, F486, and S477 via unpaired nucleotides A14 and A15, consistent with previous aptamer mutagenesis studies (Sun et al. 2021) (Fig. 3c). At the N-terminal region, C6 likely interacts with N501, a residue previously implicated in hydrogen bonding with hACE2 residues (Yan et al. 2020). While the resolution in this region limits atomic modeling of nucleotide-residue interactions, we propose aptamer binding occurs through a combination of hydrogen bonding between polar or charged amino acids (e.g., N487 and S477) and unpaired nucleotides, along with π-stacking interactions between unpaired bases and aromatic residues (e.g., F486) (Fig. 3c). Overall, these forces likely strengthen binding affinity and maintain specificity of RBD-aptamer bridge contacts, even as the complex changes position.

## 4. Discussion

In this study, we demonstrate that repurposing therapeutic aptamers provides a rapid and effective strategy for designing biosensors to detect the SARS-CoV-2 S protein. Our electrochemical aptasensor achieved high sensitivity and specificity for the SARS-CoV-2 S protein’s receptor-binding domain (RBD) within a biologically relevant detection range (Wrobel 2023; Zhu et al. 2020), demonstrating its potential for rapid and scalable diagnostic applications. We used cryo-EM to further reveal the dynamics and specificity of our aptamer interaction with the open conformation S protein. Notably, our study is the first to characterize low-resolution aptamer binding to the RBD providing structural insights into aptamer-target interactions.

The up RBD in the open conformation undergoes dynamic positional changes that influence aptamer binding. Our flexibility analysis for the up RBD aligns with prior molecular dynamics (MD) simulations and 3D variability models, which indicate that the up RBD can occupy a variety of different positions in the open conformation (Ye et al. 2022). Given that our aptamer is significantly smaller than hACE2, it is reasonable to expect a greater number of binding conformations for the aptamer-RBD interaction. In the presence of hACE2, the S protein RBDs can adopt different positions that facilitate hACE2 binding and subsequent stabilization (Yajima et al. 2024). Similarly, our aptamer-RBD complex has a stable interaction across different flexible states of the up RBD. Additionally, as the RBD transitions from a straight to a bent conformation, the sickle-shaped density between the RBDs becomes less pronounced, likely due to the up RBD moving closer to the adjacent RBD in neighboring monomers. Interestingly, this density is not present in the closed confirmation when all three RBDs are in the down configuration. We suggest this density to be a poorly resolved potential secondary aptamer-binding site in the open confirmation, as it roughly aligns with our aptamer model and localizes near the aptamer-S protein-binding residues of the two down RBDs of adjacent monomers. In conclusion, the dynamic flexibility of the up RBD allows for multiple aptamer binding conformations, and our findings highlight the stability of the aptamer-RBD interaction across different RBD positions, paralleling the behavior observed with hACE2 binding.

The visible connection between the aptamer and RBD densities allowed us to pose several hypothetical models of interaction between the aptamer and the RBD. Our C-terminal predictions are consistent with prior mutagenesis studies, highlighting the role of specific residues -particularly F486 -in replicating established hACE2 interactions (Yan et al. 2020). While previous MD simulations predicted the aptamer A17 as part of this C-terminal interface, our density-guided simulations indicate that A17 does not contribute to the A14/A15 RBD interface and may instead play a further structural role in coordinating with nearby residues or nucleotides. Our RBD N-terminal interactions provide further support for residues T500 and N501 in mimicking hACE2 binding to the aptamer (Boon et al. 2021). Given that key residues along the C- and N-terminal regions of the RBD involved in aptamer recognition also play a role in recognizing hACE2, our aptamer emerges as a strong candidate for competitively inhibiting the S protein RBD, with the potential to displace hACE2 in vivo, as demonstrated by other studies (Sun et al. 2021).

While our work highlights the sensitivity and detection mechanism of the aptasensor, further research is necessary to fully harness the potential of DNA-based biosensors for future pandemics. A key advantage of this technology is its regeneration capability, which could significantly reduce operational costs and improve scalability (Goode et al. 2015). Additionally, the minimal sample requirement of the aptasensor makes it particularly suitable for resource-limited settings, supporting rapid, minimally invasive diagnostics that are crucial for scalable outbreak responses (Broughton et al., 2020). Another advantage of the aptasensor is in its dose-dependent detection, which enhances clinical utility by enabling accurate tracking of a patient’s condition through the detection of S protein at biologically relevant concentrations. Expanding this approach, the aptamer could be adapted to target biomarkers such as acute-phase reactants and immunoglobulins, allowing clinicians to monitor disease progression and treatment efficacy, particularly in cases with fluctuating viral loads (Shanmugaraj et al., 2020). Beyond its diagnostic applications, the aptamer may also modulate S protein accessibility, potentially influencing viral entry and immune recognition. Future studies should focus on quantifying its binding kinetics relative to hACE2 and assessing its impact on S protein behavior under physiological conditions to further explore its therapeutic potential.

## 5. Conclusion

In summary, we have developed a DNA-based aptasensor for the detection of the SARS-CoV-2 S protein, demonstrating high sensitivity, specificity, and reusability. The aptasensor achieves detection within one hour, operating within biologically relevant S protein concentrations, underscoring its potential for real-time diagnostics. Our structural analysis provides insights into aptamer-RBD interactions, revealing a bridge-like binding mechanism and the flexible nature of the aptamer-RBD complex. While further optimization and clinical validation are needed, the combination of sensitivity, specificity, and scalability establishes aptasensors as promising tools for next-generation diagnostics and pandemic preparedness.

## CRediT author contribution statement

Yujun Liu: Conceptualization, Data curation, Formal analysis, Investigation, Visualization, Writing – original draft, Writing – review & editing. Kaidong Wang: Conceptualization, Data curation, Formal analysis, Investigation, Visualization, Writing – original draft, Writing – review & editing. Weiguang Wang: Formal analysis, Visualization, Writing – original draft, Writing – review & editing. Saarang Kashyap: Formal analysis, Visualization, Writing – original draft, Writing – review & editing. Jonathan Jih: Formal analysis, Writing – review & editing. Anthony Imani: Formal analysis, Visualization, Writing – original draft, Writing – review & editing. Tzung K. Hsiai: Conceptualization, Methodology, Supervision, Writing – review & editing, Funding acquisition, Resources. Z. Hong Zhou: Conceptualization, Methodology, Project administration, Supervision, Writing – original draft, Writing – review & editing, Funding acquisition, Resources. All authors reviewed and approved of the paper.

## Declaration of competing interest

The authors declare no competing interests.

## Acknowledgments

We acknowledge use of resources in the Electron Imaging Center for Nanomachines supported by UCLA and grants from the NIH (1S10OD018111) and the National Science Foundation (DBI-1338135 and DMR-1548924). Molecular graphics and analyses performed with UCSF ChimeraX, developed by the Resource for Biocomputing, Visualization, and Informatics at the University of California, San Francisco, with support from National Institutes of Health R01-GM129325 and the Office of Cyber Infrastructure and Computational Biology, National Institute of Allergy and Infectious Diseases.

## Funding

This work was supported by grants from the National Institutes of Health (NIH) to T.K.H. (R01HL149808, R01HL118650, R01HL159970, R01HL129727, and T32HL144449) and to Z.H.Z. (R01GM071940 and R01DE025567), and from VA I01BX004356 (T.K.H.).

## Additional information

All data needed to evaluate the conclusions in the paper are present in the paper and/or the Supplementary Information. Additional data related to this paper may be requested from the authors. Cryo-EM density maps and 3D models are deposited in EMDB and PDB under the accession numbers EMD-XXXXX and XXXX, respectively.

## Notes

### Competing Interest Statement

The authors have declared no competing interest.

