## Supplemental Table 1 for "Demonstration and structural basis of a therapeutic DNA aptamer for SARS-CoV-2 spike protein detection"

**Supplementary Table S1. CryoEM data collection**

| | Open Conformation C1 (EMD-XXXXX) | Closed Confirmation C1 (EMD-XXXXX) | Closed Confirmation C3 (EMD-XXXXX) |
|---|---|---|---|
| **Data collection and processing** | | | |
| Magnification | 81,000 | 81,000 | 81,000 |
| Voltage (kV) | 300 | 300 | 300 |
| Electron exposure (e–/Å$^2$) | 44 | 44 | 44 |
| Defocus range (μm) | -1.5 to -2.5 | -1.5 to -2.5 | -1.5 to -2.5 |
| Pixel size (Å) | 1.1 | 1.1 | 1.1 |
| Symmetry imposed | C1 | C1 | C1 |
| Particle images (no.) | 39,240 | 141,547 | 141,547 |
| Map resolution (Å) | 4.4 | 3.8 | 3.4 |
| FSC threshold | 0.143 | 0.143 | 0.143 |
| Symmetry imposed | C1 | C1 | C3 |
